# PanIN and CAF Transitions in Pancreatic Carcinogenesis Revealed with Spatial Data Integration

**DOI:** 10.1101/2022.07.16.500312

**Authors:** Alexander T.F. Bell, Jacob T. Mitchell, Ashley L. Kiemen, Kohei Fujikura, Helen Fedor, Bonnie Gambichler, Atul Deshpande, Pei-Hsun Wu, Dimitri N. Sidiropoulos, Rossin Erbe, Jacob Stern, Rena Chan, Stephen Williams, James M. Chell, Jacquelyn W. Zimmerman, Denis Wirtz, Elizabeth M. Jaffee, Laura D. Wood, Elana J. Fertig, Luciane T. Kagohara

## Abstract

Spatial transcriptomics (ST) is a powerful new approach to characterize the cellular and molecular architecture of the tumor microenvironment. Previous single-cell RNA-sequencing (scRNA-seq) studies of pancreatic ductal adenocarcinoma (PDAC) have revealed a complex immunosuppressive environment characterized by numerous cancer associated fibroblasts (CAFs) subtypes that contributes to poor outcomes. Nonetheless, the evolutionary processes yielding that microenvironment remain unknown. Pancreatic intraepithelial neoplasia (PanIN) is a premalignant lesion with potential to develop into PDAC, but the formalin-fixed and paraffin-embedded (FFPE) specimens required for PanIN diagnosis preclude scRNA-seq profiling. We developed a new experimental pipeline for FFPE ST analysis of PanINs that preserves clinical specimens for diagnosis. We further developed novel multi-omics analysis methods for threefold integration of imaging, ST, and scRNA-seq data to analyze the premalignant microenvironment. The integration of ST and imaging enables automated cell type annotation of ST spots at a single-cell resolution, enabling spot selection and deconvolution for unique cellular components of the tumor microenvironment (TME). Overall, this approach demonstrates that PanINs are surrounded by the same subtypes of CAFs present in invasive PDACs, and that the PanIN lesions are predominantly of the classical PDAC subtype. Moreover, this new experimental and computational protocol for ST analysis suggests a biological model in which CAF-PanIN interactions promote inflammatory signaling in neoplastic cells which transitions to proliferative signaling as PanINs progress to PDAC.

**Summary:** Pancreatic intraepithelial neoplasia (PanINs) are pre-malignant lesions that progress into pancreatic ductal adenocarcinoma (PDAC). Recent advances in single-cell technologies have allowed for detailed insights into the molecular and cellular processes of PDAC. However, human PanINs are stored as formalin-fixed and paraffin-embedded (FFPE) specimens limiting similar profiling of human carcinogenesis. Here, we describe a new analysis protocol that enables spatial transcriptomics (ST) analysis of PanINs while preserving the FFPE blocks required for clinical assessment. The matched H&E imaging for the ST data enables novel machine learning approaches to automate cell type annotations at a single-cell resolution and isolate neoplastic regions on the tissue. Transcriptional profiles of these annotated cells enable further refinement of imaging-based cellular annotations, showing that PanINs are predominatly of the classical subtype and surrounded by PDAC cancer associated fibroblast (CAF) subtypes. Applying transfer learning to integrate ST PanIN data with PDAC scRNA-seq data enables the analysis of cellular and molecular progression from PanINs to PDAC. This analysis identified a transition between inflammatory signaling induced by CAFs and proliferative signaling in PanIN cells as they become invasive cancers. Altogether, this integration of imaging, ST, and scRNA-seq data provides an experimental and computational approach for the analysis of cancer development and progression.

## Main text

Single-cell RNA-sequencing (scRNA-seq) and spatial molecular technologies have enabled unprecedented characterization of the molecular and cellular pathways that comprise the tumor microenvironment (TME) and its function pan-cancer^1,2^. This has had a particularly profound impact on the understanding of the complex immunosuppressive environment that characterizes pancreatic ductal adenocarcinoma (PDAC) and its hypothesized role hypothesized role in cancer progression^3–7^. However, further characterization of premalignancy is needed to delineate the precise evolutionary mechanisms that underlie malignant transformations and to understand the impact of the complex microenvironment that facilitates carcinogenesis and the development of nearly universal therapeutic resistance in PDAC. Pancreatic intraepithelial neoplasia (PanIN) is a pre-malignant lesion that progresses into pancreatic ductal adenocarcinoma (PDAC), and thereby provides the opportunity to directly characterize these evolutionary processes. However, the diagnosis of preinvasive cancer lesions, such as PanINs, is almost always limited to formalin-fixed and paraffin embedded (FFPE) tissue and most of the current knowledge on the biology of these lesions is restricted to bulk transcriptional analysis.

Thus, the recent development of a spatial transcriptomics (ST) technology that can utilize FFPE samples will provide untapped opportunities to apply high dimensional approaches to evaluate archived FFPE specimens of preinvasive PanIN biospecimens, with broad applicability to further biobanks of clinical trials as well as standard of care diagnostic samples^8^. Spatial molecular technologies can drive pathway discoveries in cancers and their TME while preserving tissue architecture, enabling the characterization of molecular changes that result from cell-to-cell direct interactions^2^. ST approaches have already identified transcriptional signatures associated with spatial interactions that are delineating the cellular phenotypes that underlie tumor biology, evolution, and responses to therapy^2^. Until recently, these approaches were limited to fresh frozen sample profiling that maintain RNA integrity^9–13^. The newly developed ST approach for FFPE will leverage the accessibility to major sources of biopsy and surgical tissue samples stored in paraffin blocks and will allow retrospective studies spanning stages of cancer evolution.

An additional challenge regarding the use of FFPE samples is the section preparation as the platform limits the size of the tissue area that can be analyzed. Extensions of ST technology which enable direct profiling on custom slides surmount this limitation. However, sectioning of these valuable FFPE biospecimens for ST analysis can destroy their integrity for subsequent clinical diagnostic testing, which must take precedent to single-cell analysis in translational research studies. Therefore, this study develops new methods to prepare smaller sections from FFPE blocks for ST analysis that uniquely preserves limited and valuable clinical samples, including PanINs.

Computational analysis methods to discovery the cellular and molecular changes in the evolution of the microenvironment are an essential component of ST technologies, and are being actively developed alongside the experimental approach^2^. Multi-omics analysis, including integration with scRNA-seq data, is of particular interest for both deconvolution methods that overcome the lack of single-cell resolution of spatial technologies^14^ and contextualization of pathways learned about the TME within the context of larger scRNA-seq reference atlases. The combination of imaging with ST data enables a further level of integration to improve analysis, and are currently being developed to augment clustering of ST data for cell type annotation^15,16^. In FFPE ST, the possibility of staining and imaging the sections prior to library preparation allows pathological examination of cell morphologies to enhance annotations of cellular function at a single-cell resolution that are disregarded in current integration methods. We recently developed a machine learning method that provides 3-dimensional (3D) pathologic tissue assessments, named CODA, to identify and quantify normal and PanIN cells in the pancreas from hematoxylin and eosin (H&E) stained FFPE sections^17^. This advance combined with our recent collation of a robust single-cell atlas of 61 PDAC tumors provides the unique opportunity for an innovative, new three-fold integration of imaging, ST, and scRNA-seq data that is tailored to analyzing the dynamic cellular transitions throughout PDAC initiation and progression to advanced disease.

In this study, we develop and apply novel experimental and computational approaches for ST analysis to a cohort of PanINs spanning low-grade and high-grade lesions. Analysis of the epithelial cells purified through integration with imaging using CODA revealed that all except one of the PanIN lesions share similar expression patterns to the classical PDAC subtype. The one PanIN sample that does not express this classical signature expresses cancer stem cell (CSC) markers, suggesting that cells with stemness transcriptional features are present at premalignant stages. Moreover, this integrative analysis demonstrated that the PanINs microenvironment contains the same cancer associated fibroblast (CAF) subtypes that are enriched in invasive PDACs, providing new evidence that these cells modulate premalingnacy. The further multi-omics integration with our reference scRNA-seq atlas of PDAC further found that a CAF-associated inflammatory pattern in neoplastic epithelial cells gradually decreases during PDAC invasion, and is associated with a compensatory increase in proliferation pathways in PDAC carcinogenesis. Altogether, this integrated experimental and computational approach for multi-omics integration is broadly applicable to analysis of cancer progression in other tumor types.

### Spatial transcriptomics applied to FFPE specimens captures preneoplastic pancreatic tissue architecture

To identify the mechanisms associated with PanIN to PDAC progression, we applied a recently developed FFPE ST protocol to PanIN samples and the analysis combined two transfer learning methods (**Figure 1A**). The ST slide’s dedicated areas for analysis are small (6×6 mm), and the FFPE preserved samples are typically larger and would require coring or scraping of the block to isolate only the PanIN lesion. Extensive manipulation of FFPE blocks for the ST sample preparation would compromise future clinical diagnostics assessment of the broader PDAC surgical specimen associated with the PanIN lesions selected. Therefore, we developed a method to score the surface of the FFPE blocks using 5mm diameter circular skin biopsy punches. Prior to sectioning, the punches were used to score the regions containing the ~1mm PanIN lesion while preserving the block. The scoring then allowed the non-relevant tissue to detach from the regions that were collected and placed on the designated area of the ST slide (**Figure 1A**). This approach enables ST profiling of selected PanIN lesions, while minimizing manipulation to protect the FFPE block for further analysis. Following sections preparation and ST data generation, we incorporated a machine learning approach, CODA, to identify cell types in each sample and at the same time deconvolute the cell types captured in each ST spot (**Figure 1A**). The accurate cell classification for each ST spot allowed more robust differential analysis between normal and PanIN spots. To integrate the PanIN analysis with PDAC samples to build the carcinogenesis process of pancreatic cancer, we applied a transfer learning method, ProjectR, to identify PanIN signatures in a scRNA-seq PDAC compilation dataset, and vice-versa (**Figure 1A**).

**FIGURE 1.**
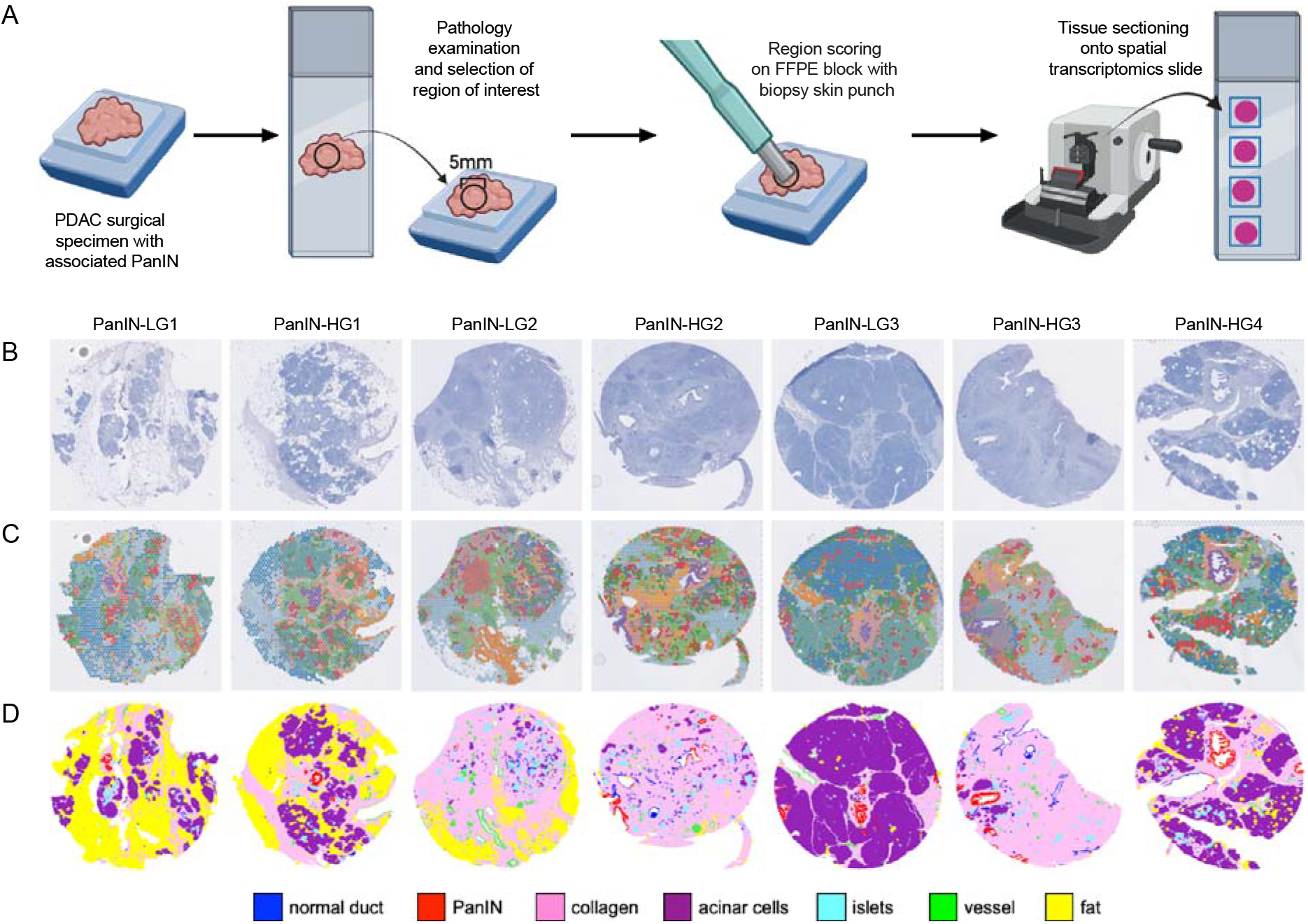
Spatial transcriptomics analysis of FFPE pancreatic intraepithelial neoplasia (PanIN). (A) Pancreatic cancer surgical specimens in FFPE were examined and the regions containing PanIN lesions were identified for scoring using a 5mm skin biopsy punch and sectioning onto the spatial transcriptomics slide. (B) Stained sections were used for pathology examination and identification of PanINs and other pancreatic histological regions. (C) The unsupervised clustering of the spatial transcriptomics data identified gene expression clusters which location resembles the distribution observed in the stained sections. (D) Single-cell resolution of the cell subtypes indicated in the legend were defined automatically from cellular morphologies of the H&E imaging using the machine learning approach CODA^16^, thereby refining cellular annotations obtained from clustering alone.

To study the mechanisms of progression from pre-malignant early PanINs to PDAC, we applied this FFPE ST protocol to profile 4 patient tumor specimens with paired low- (LG) and highgrade (HG) PanINs (total number of lesions = 8). This cohort was designed to enable comparisons of progressive mechanisms within and between patients. Initial total RNA quality check indicated that all samples presented some level of RNA degradation (RIN ~2) but with a high concentration of 200bp fragments (DV200 > 50%) compatible with the FFPE ST platform. Following ST data generation, pre-processing and filtering, seven out of the eight samples presented high quality data for subsequent downstream analysis. We were able to detect an average of 71,695 reads and 2,537 genes per spatial spot, and an average of 16,266 genes per sample.

The ST data from our PanIN cohort provides combined image and transcriptomics profiling (**Figures 1B and 1C**). We characterized the canonical cellular distribution of PanINs and surrounding pancreas tissue by first applying clustering to the ST profiling data alone. The normal pancreas is composed of multiple cell subtypes with different functions. To execute its most important functions, the pancreas is composed of exocrine cells (acinar cells) that are responsible for the production of digestive enzymes and by endocrine cells (islets of Langerhans) that produce insulin and other hormones. The excretion of enzymes occurs through pancreatic ducts, while insulin and hormones are directly released into the blood stream. PanINs and PDACs differentiation resembles the morphology of normal pancreatic ducts^19^. Annotating marker genes differentially expressed in each cluster learned from the transcriptional signal infers these canonical and transformed cell types (**Figure 1C, Supplemental Figure 1**). Moreover, the distribution of the normal and neoplastic cell types identified from clustering matches their locations in the corresponding section image (**Figures 1B and 1C**). In all samples, it was possible to map specific clusters to the PanINs that were distinct from the other clusters including the normal ducts. Nevertheless, in some regions the clusters extend beyond cellular boundaries into the adjacent cell types on H&E imaging (**Supplemental Figure 2**). This extended signal could arise either from the intercellular signaling extending the molecular changes beyond cellular margins or from technical artifacts in the ST technology.

To improve cluster annotations based on tissue morphology, we applied a machine learning method, named CODA, to the H&E data from Visium to automatically classify the pancreatic normal and neoplastic cells at a single-cell resolution to enhance the spot-based ST analysis. Briefly, CODA is a 3D imaging-based approach that uses deep learning semantic segmentation to identify different cell types within the human pancreas (acinar cells, islets of Langerhans, fibroblasts, adipocytes, endothelial cells, ductal cells, neoplastic cells)^17^. In this study, we adapted CODA for integration of imaging with Visium to obtain a color-coded image with each color corresponding to a specific cell type from the stained tissue sections (**Figure 1D**). In contrast to the clustering annotations, cells annotated through imaging using CODA are at singlecell resolution. Thus, we could apply this approach to estimate the true proportion of cells within each spot (**Supplemental Figure 3**) associated with each spatial barcode in the ST data for further downstream analysis. By determining cell proportions in each ST spot we were able to select those representing a unique cell type, while filtering out spots capturing multiple cell types, to avoid unwanted bias in the comparisons between normal and PanIN clusters, for example.

### PanIN-associated fibroblasts are a heterogeneous population composed of the same subtypes detected in invasive PDAC

The ability of our integration of imaging and ST data to characterize PanINs and their surrounding microenvironments provides the unique opportunity to examine the fibroblast population adjacent to these lesions. While CODA broadly annotates stromal cells (**Figure 2A**), the PDAC TME is enriched with a heterogeneous population of cancer associated fibroblasts (CAFs). They have been classified into three subtypes based on transcriptional profiles: myofibroblastic CAFs (myCAFs), inflammatory CAFs (iCAFs), and antigen-presenting CAFs (apCAFs)^21–23^. These populations of mesenchymal cells play dual roles and can induce or inhibit PDAC progression^21–24^. CAFs exert a tumorigenic role by providing metabolites for tumor cell survival, stimulating cell growth pathways through paracrine signaling, and creating an immunosuppressive microenvironment^22^. However, a tumor suppressing CAF enriched TME can reduce essential nutrients required for tumor progression and differentiation, and the same CAFs can be functionally repolarized to release chemokines that will recruit immune cells into the tumor^22^.

**FIGURE 2.**
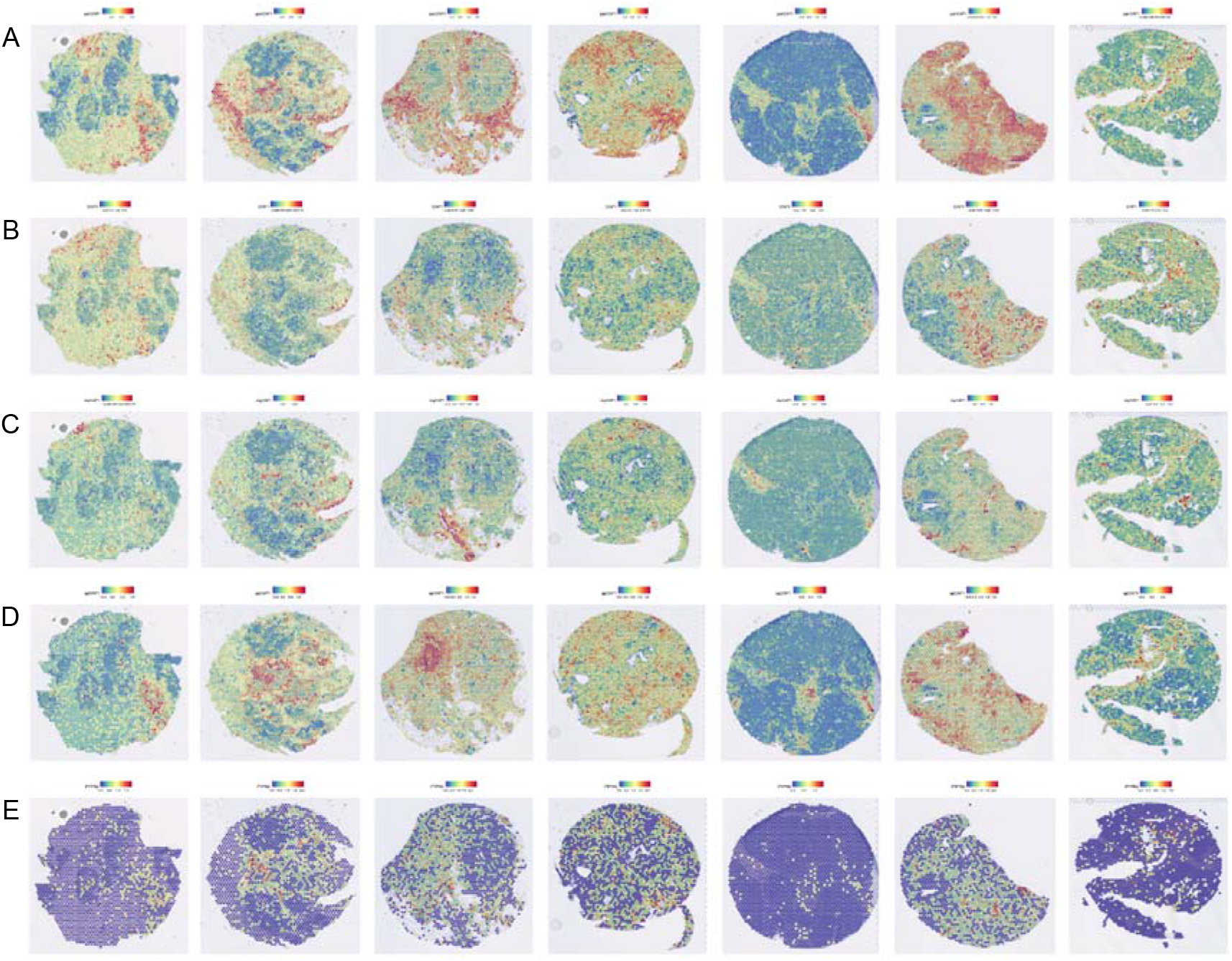
Spatial distribution of PDAC cancer associated fibroblasts (CAF) subtypes. (A) CAFs localization was mapped using pan-CAF markers, (B) myofibroblastic-CAF markers, (C) inflammatory-CAF markers and (D) antigen presenting-CAF markers. (E) CD45 expression was examined to identify regions where CAFs and immune cells were co-localized.

MyCAFs and iCAFs have previously been observed in pancreatic premalignant lesions in murine models that recapitulate PDAC development, suggesting that they arise early during tumorigenesis^25,26^. Nevertheless, their presence is not well described in human premalignant lesions. Here, we leveraged our computational analysis approach to purify stromal cells in the ST data and further classify these cells from the ST data using established gene markers^23^ to map the distribution of myCAFs, iCAFs and apCAFs in the human PanINs microenvironment. In our cohort, the density of stromal cells inferred from CODA varied but were observed adjacent to each premalignant lesion (Figure 1D, pink annotated regions). The further integration of CODA annotations with the ST transcriptional profiles showed that a CAF common signature (pan-CAF) is consistently expressed across the collagen rich regions annotated by CODA (**Figure 2B**, orange and red spots). The expression of myCAF (**Figure 2C**) and iCAF (**Figure 2D**) markers were detected in all samples overlapping with the regions where pan-CAFs are present.

The presence of a recently described subtype of apCAFs was also investigated using the transcriptional data in our cohort. apCAFs were first identified by scRNA-seq in a PDAC mouse model. Further characterization showed that these cells express MHC-II genes and can present antigens to CD4^+^ T cells in vitro^23^. Subsequently, apCAFS have been shown to present antigen to Tregs which activates their suppressive capability^27^. In our study, expression of the apCAF signature was detected in all samples (**Figure 2E**). As expected, a significant proportion of the apCAF positive spots also express CD45, a marker of immune cells (**Figure 2F**) that can express the same MHC-II genes. Since CODA cannot annotate immune cells due to their limited size and scant cytoplasm, and ST does not provide single-cell resolution, the confirmation of apCAFs in some samples could not be exactly defined by our analysis. Nevertheless, in some regions of the stroma the expression of apCAF markers do not colocalize with CD45+ cells, indicating that these mesenchymal cells are present in human PanINs.

### ST identifies expression of both PDAC classical subtype and cancer stem cell signatures in PanINs

CAFs are mediators of PDAC progression and aggressiveness through interactions with neoplastic cells^22,23,28^. The detection of PDAC associated fibroblast subtypes in PanINs prior to establishment of invasive carcinoma suggests that the differentiation of fibroblasts into CAFs is an early event that may influence PanIN progression to PDACs. To test this, we leveraged the automated cell type annotation from CODA with cluster-based annotations to select spots with >70% purity of ductal cells to compare normal and PanIN ducts (**Figure 3A**). Next, we characterized PanIN cell heterogeneity relative to the established classical and basal-like PDAC subtypes^20^. We found that six out of seven PanINs express the PDAC classical subtype signature (**Figure 3B**). The basal-like signature is not detected in any of the premalignant lesions (**Figure 3C**). This observation supports the hypothesis that PDACs arise with a classical phenotype and likely acquire the basal-like phenotype upon progression and accumulation of molecular aberrations^30^.

**FIGURE 3.**
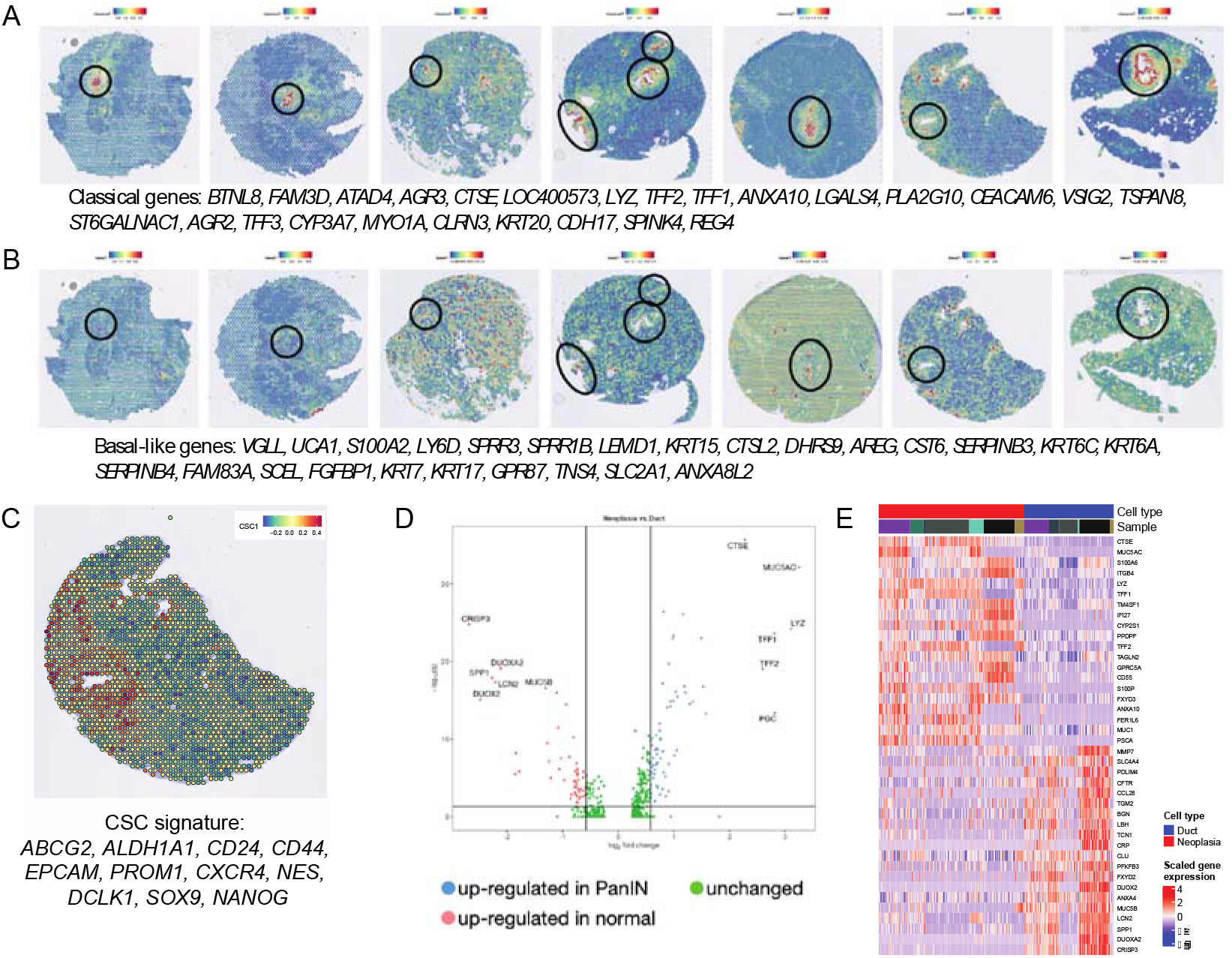
Pancreatic intraepithelial neoplasia (PanINs) transcriptional features. (A) Six out of seven PanINs, expressed markers that characterize the classical subtype of pancreatic cancer, while (B) the basal-like signature was not expressed by any of the premalignant lesions. (C) The only sample that is neither classical nor basal-like expresses cancer stem cell (CSC) markers. (D) Differential expression analysis identified genes which up-regulation (blue dots) or downregulation (red dots) in PanINs, relative to normal ducts, discriminate preneoplastic from normal cells (E).

Only one HG PanIN sample (PanIN-HG3, Figure 1) expressed neither the classical nor the basal-like signatures. Thus, we hypothesized that this sample expresses a third transcriptional phenotype. PDAC progression, resistance to therapies, and immune evasion are partialy associated with the presence of PDAC cells expressing cancer stem cell (CSC) markers21. We verified the expression of CSC markers among the PanINs in the cohort. The only sample with significantly high expression of CSC markers is the one that did not express the classical or the basal-like PDAC signatures (**Figure 3D**). The presence of cells expressing CSC markers in PanINs was previously described in a mouse model that mimics PDAC development^22^ and in human samples^23^, but little is known about the mechanisms leading to CSC genes up-regulation their role in PDAC progression. Our observation that this stemness signature is not observed in cells expressing the classical subtype suggests that neoplastic cells with stemness features are a distinct population that arise in early pre-malignant stages. Nonetheless, further investigation in a larger cohort is needed to determine the frequency of this stem-cell mechanism of progression, the pathways driven by stemness, and how these cells are interacting with the CAFs and other cells in the TME to modulate PDAC biology.

### Differential expression analysis between PanINs and normal ducts identifies gradual *TFF1* increased expression during PanIN progression limited to the classical phenotype

To further define the molecular features of PanINs, spots from all samples CODA annotated as normal and PanIN ducts were merged for each patient and differential expression was performed to identify gene expression changes across each patient’s premalignant lesions. A total of 118 genes are differentially expressed in PanINs relative to normal ducts (**Figure 3E**) and their expression pattern discriminated PanINs from normal ducts among the different samples (**Figure 3F**). Among the top 20 up-regulated genes in the premalignant lesions, only 5 genes (*TM4SF1, CYP2S1, CD55, FER1L6* and *PSCA*) had no known role in pancreas tumorigenesis, suggesting that FFPE ST analysis is robust and corroborates previous gene expression analyses in PanINs^34–36^. The pathway analysis from the differentially expressed genes indicates enrichment for MYC and oxidative phosphorylation pathway mediators. Both signaling pathways have been previously shown to be upregulated in PanINs and PDAC, particulary in association with progression from premalignancy to invasive cancer, metastasis development, and resistance to therapy^24–26^ (**Supplemental Figure 4**). Although predominantly consisting of the classical subtype, the differential expression analysis highlighted the inter-sample heterogeneity with only one differentially expressed gene showing up-regulation in all classical samples (*TFF1*). *TFF1* is known to be up-regulated in PanINs and PDACs and its protein levels have been suggested as a potential early detection marker found in bodily fluids. In in vitro cell culture models, the secreted form of TFF1 has been shown to increase PDAC and stellate cell motility without a significant impact on proliferation^27^. Since stellate cells are considered one of the precursors to some PDAC CAF subtypes^28,29^, it is possible that *TFF1* is one of the mediators of intercellular interactions between PanIN and PDAC cells and CAFs. However, the sample expressing the CSC markers signature does not express *TFF1*, suggesting that the stemness signature and *TFF1* are mutually exclusive (**Supplemental Figure 5**).

The characterization of multiple ducts, including those spanning across stages of PanIN differentiation (mixed ducts), provides the opportunity to trace the cellular changes associated with PanIN progression. Additionally, ST analysis provides the ability to visualize the preneoplastic differentiation stages and concomitantly map the respective gene expression level changes (**Figure 4A**). We therefore compared expression changes between lesions classified as LG or HG based on their morphology. The differentiation stages of PanINs cannot be discriminated using CODA and the classification of LG and HG preneoplastic ducts was performed through pathology examination (KF and LDW) (**Figure 4B,C** and **D**). Using the pathological PanIN classification, we identified mixed ducts containing normal, LG and HG preneoplastic cells (**Figure 4E**).

**FIGURE 4.**
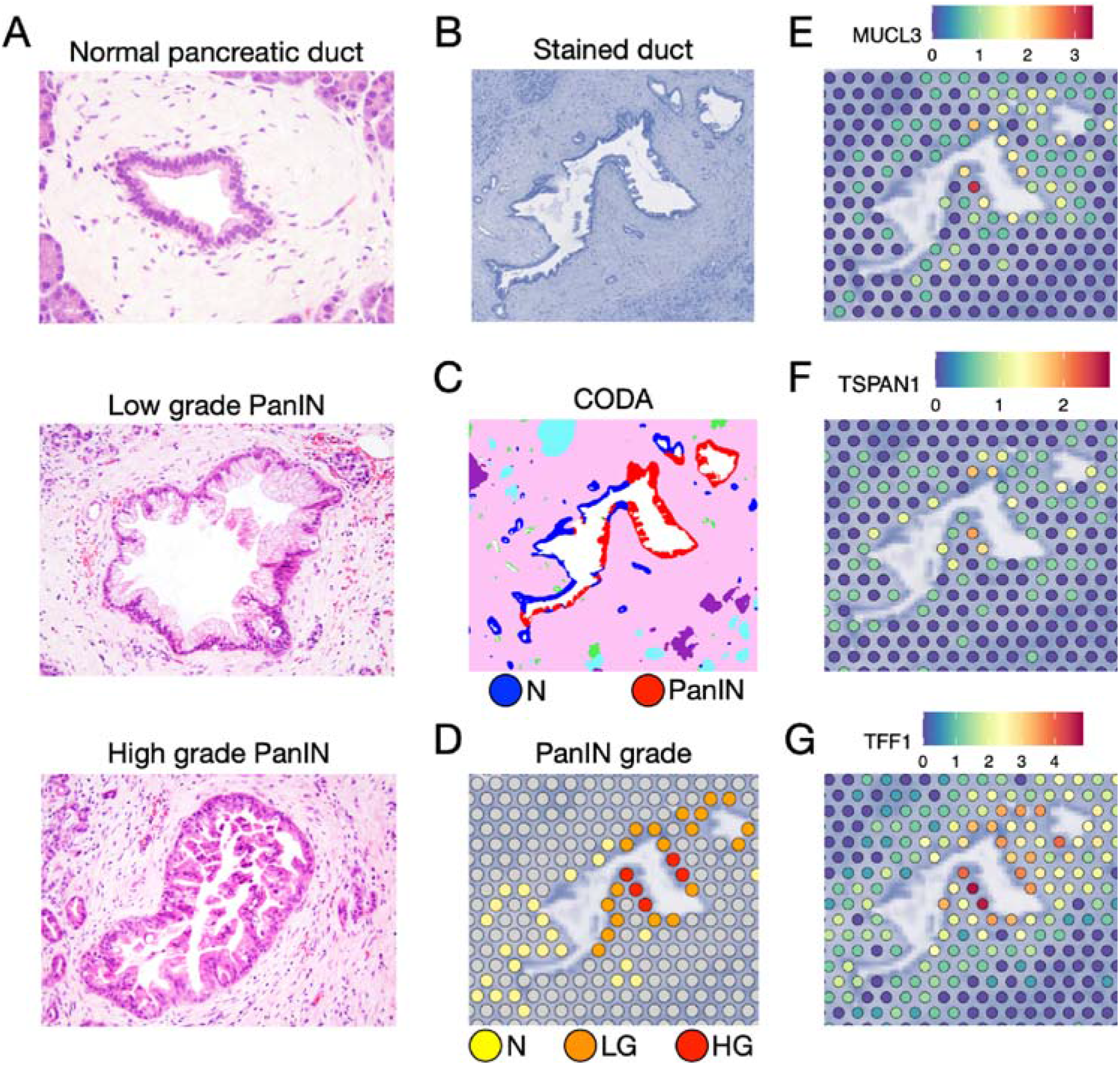
Identification of transcriptional changes associated with pancreatic intraepithelial neoplasia (PanIN) differentiation grade. (A) Normal, low grade (LG) and high grade (HG) PanINs are morphologically distinct and can be classified by pathology examination. (B, C and D) As a model for PanIN progression, a mixed pancreatic duct containing normal, LG and HG cells was used to better visualize changes in expression. Top genes from the differential expression analysis, (E) *MUCL3*, (F) *TSPAN1* and (G) *TFF1*, show gradual increase from normal through LG until HG progression.

We expanded our differential expression analysis study to uncover additional gene expression changes across PanIN stages. This analysis identified five other genes (*MUCL3, C19orf33, TSPAN1, SCD*, and *ACTB*) that were up-regulated in HG lesions relative to LG lesions (**Supplemental Figure 7**). In addition, the level of expression of *MUCL3* and *TSPAN1*gradually increased from normal ducts through LG and HG lesions (**Figure 4F** and **4G**). The same pattern was observed for *TFF1* which was found to be up-regulated in the PanIN expressing the classical PDAC genes. This gradual change in expression is best visualized in one of the PanIN samples in which a single duct presents a mix of normal, LG and HG cells (**Figure 4H**).

### Changes in PanIN progression map to transitions in malignancy in PDAC

PanINs are premalignant lesions that can progress to PDAC; this is supported by the detection of common driver mutations in premalignancies that are frequent in invasive cancer^30–32^. The examination of other molecular alterations that are present in PanINs and conserved in PDACs could provide new knowledge about the early transcriptional events of pancreatic carcinogenesis and the mechanisms driving the continuous development into invasive cancer. A limitation of our cohort in making these inferences is its small sample size and profiling only the PanIN lesions across different individuals, and the lack of similar profiling of PDAC tumors from those patients. Our combined set of public domain scRNA-seq PDAC datasets (PDAC Atlas)^18^ provides a cohort of over 61 samples against which we can further validate the molecular changes observed across grades of differentiation. While these single-cell datasets lack the imaging resolution of their ST counterparts, the dissociation procedures applied to the pancreas would be anticipated to retain cells from PanIN lesions when they co-occur with tumors. Therefore, further computational methods for multi-omics integration of our ST data of PanINs and scRNA-seq data in the PDAC atlas can further our analysis of molecular changes in the neoplastic epithelial cells that underly carcinogenesis (**Figure 5A**).

**FIGURE 5.**
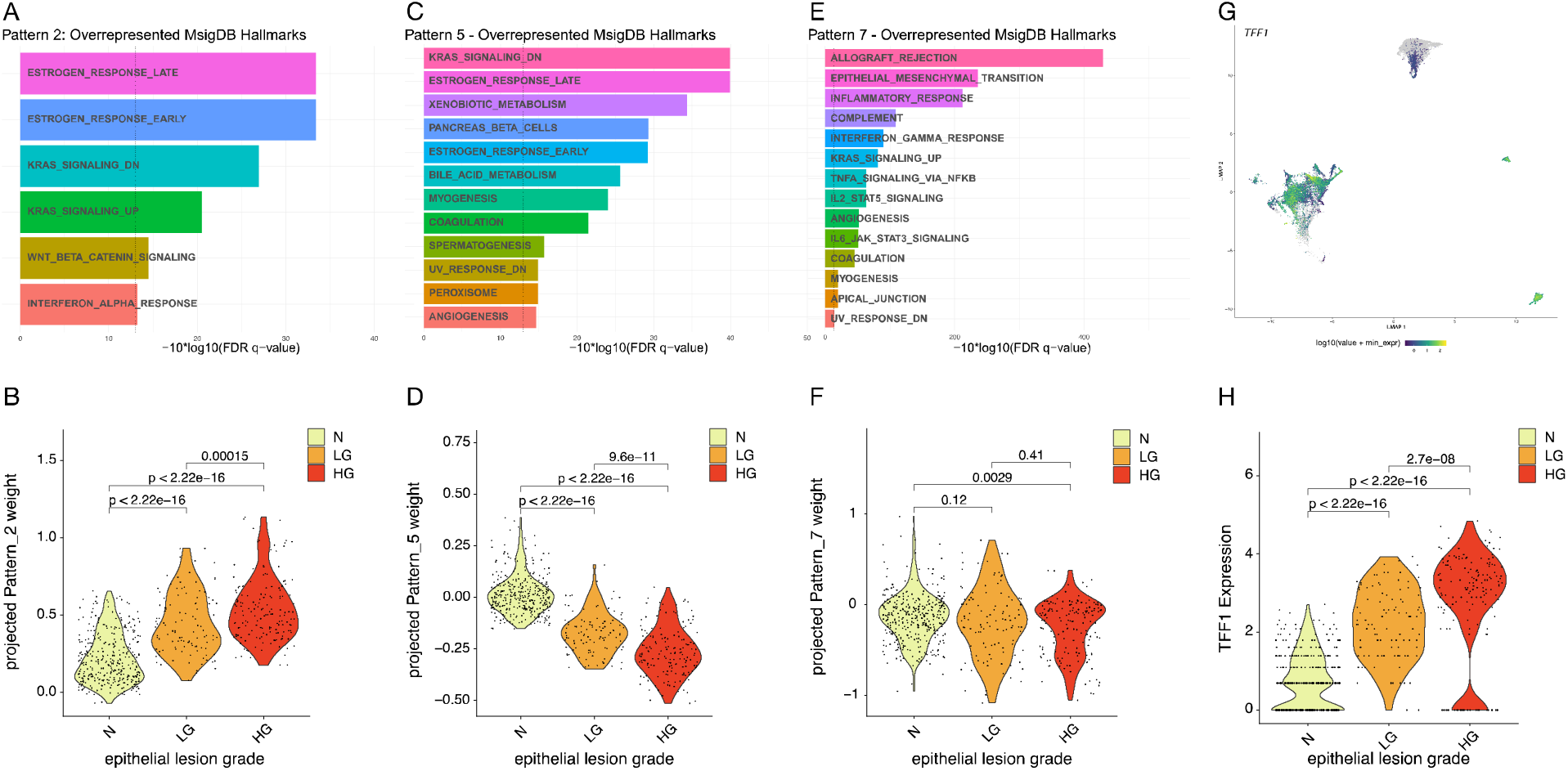
Integration of pancreatic intraepithelial neoplasia (PanIN) spatial transcriptomics (ST) data with invasive pancreatic cancer single-cell RNA-sequencing (scRNAseq) using transfer learning. (A, C and E) Representation of enriched MSigDB pathways in Pattern 2, Pattern 5 and Pattern 7 of the PDAC atlas. (B, D and F) Violin plots of projected PDAC atlas Pattern 2, Pattern 5 and Pattern 7 weights in PanIN ST spots. (G) UMAP embedding of epithelial cells from the PDAC atlas colored by *TFF1* expression. (H) Violin plots of TFF1 expression in all PanIN ST epithelial spots grouped by epithelial lesion grade. P-values were calculated using two-sample Wilcoxon rank-sum tests. (N: normal, LG: low-grade, HG: high-grade).

To enable this analysis of tumor progression, we selected the gene expression profiles of 25,442 epithelial ductal cells in 61 biospecimens collated from six previously published scRNA-seq datasets (Cancer: 14,589 cells; Normal: 7,561 cells; Normal Tumor Adjacent: 2,375 cells, Unspecified: 917 cells) (**Supplemental Figure 8A**). UMAP analysis of these cells identifies a phenotypic switch between true normal epitheial cells, tumor-adjacent normal cells, and within malignant epithelial cells supporting our hypothesis that these datasets likely contain unannotated PanIN cells. Using the PDAC atlas, we verified that *TFF1* expression increases between normal epithelial cells and in tumor-adjacent normal epithelial cells and then again further increases in a subset of malignant PDAC cells (**Figure 5B**), mirroring the stage-specific increase in its expression observed in PanIN cells (**Figure 5C**). This integrative analysis further supports the association of this gene with PanIN and invasive PDAC progression*. TFF1* expression is almost undetectable in normal ductal cells. Surprisingly, the normal ductal cells adjacent to tumor cells express low levels of *TFF1*, suggesting that the transcriptionally normal surrounding ducts are already programmed toward a pre-malignant state.

To further delineate the molecular transitions that the malignant epithelial cells cells undergo, our complementary study of the PDAC atlas applied the Bayesian non-negative matrix factorization method CoGAPS^48,49^ that learned eight transcriptional patterns that delineate transitions in the epithelial cells^18^. In this current study, we seek to integrate the patterns learned from the scRNA-seq data with our ST dataset to determine the extent to which they represent stage-related transitions in the transformation from PanIN to PDAC. To enable the integrative analysis between ST and scRNA-seq data, we adapted out transfer learning approach projectR^33,34^ to spatial data integration by projecting the patterns learned in the scRNA-seq data onto the epithelial spots from the ST data (N = 623 spots; normal = 254, LG = 110, HG = 159). Among the patterns projected from the atlas onto the ST data, Pattern 2 (**Figure 5D** and **Supplemental Figure 8B**), enriched with genes involved in KRAS signaling and estrogen response, showed a marked increase in projected pattern weights from normal epithelium through LG and HG PanINs (**Figure 5E** and **Supplemental Figure 8C**), corroborating previously reported studies showing up-regulation of pancreatic oncogenic signaling pathways in premalignancy initiation and progression^50,51^. Pattern 5 (**Figure 5F** and **Supplemental Figure 8D**), associated with normal metabolic activity, showed a substantial decrease in projected pattern weights with progression of PanIN lesion grade (**Figure 5G** and **Supplemental Figure 8E**), suggesting a decline of normal cellular function in the PanIN cells following the same trend that is present in the primary tumor cells from the PDAC Atlas. Pattern 7 (**Figure 5H** and **Supplemental Figure 8F**), representing an inflammatory state, is enriched in normal ductal cells and dissipates with the development of early stage PDAC and progression to advanced cancer. Pattern 7 also showed decreasing levels over the course of progression from normal cells to PanIN (independent of the differentiation grade), as demonstrated by the increase in the number of spots with low projected weights (**Figure 5I** and **Supplemental Figure 8G**). This analysis provides further evidence that the scRNA-seq data in the PDAC atlas contains unclassified PanIN cells, and yields a model in which inflammatory signaling in epithelial cells precedes further signaling changes that promote growth in malignant cells during carcinogenesis.

## Discussion

ST technologies are uncovering new molecular and intercellular interactions that provide insights into how these complex signaling networks mediate cancer development and progression^2^. In this study, we expanded a novel protocol developed for FFPE ST^8^ to preserve FFPE blocks and thereby enable the analysis of PanINs to uncover the mechanisms of progression from premalignancies to invasive PDAC. Another innovation of our study is the development of two machine learning methods for integrative analysis across imaging, ST profiles, and scRNA-seq data for the analysis of these data. The first method, CODA^17^, enabled the assignment of cell types to ST spots at a true single-cell resolution based on the imaging of the each ST section. Applying this imaging and ST integration facilitated accurate assignment of cell types, isolation of spots with single cell types for analysis, providing a framework for future ST deconvolution methods for analysis. The second method, ProjectR^33,34^, allowed the integration of scRNA-seq from invasive PDACs with ST data from PanINs to relate the mechanisms associated with PDAC initiation to subsequent PDAC progression. While our analysis applied this multi-omics approach to isolate molecular changes in purified cellular subpopulations, we note that the imaging-based cellular annotations and further scRNA-seq data can also be readily extended to spot-based deconvolution of ST data. In this study, our integration of state-of-the-art experimental and computational approaches allowed us to characterize the molecular and cellular features of PanIN to PDAC development, providing a multi-omics framework for the broader study of carcinogenesis.

Applying our combined experimental and computational approach to PanIN samples, we observed for the first time the presence of CAFs and the different transcriptional subtypes (myCAF, iCAF and apCAF) in premalignant human lesions. These subtypes were only previously described in PDAC^21,23^. CAFs are the most abundant cell type in the PDAC TME and are known to influence tumor cell behavior and to create an immunosuppressive environment^35^. The presence of these regulatory cells in human pancreatic premalignant lesions is not well described, but suggests that CAF-induced TME remodeling is an early event with durable impact on PDAC development. Further studies are necessary to examine the specific interactions driven by the different CAF subtypes and how they modulate preinvasive neoplastic cells and other cellular components of the PDAC TME. Such knowledge is critical to guide the development of new therapeutic interventions that inhibit or revert CAF oncogenic and immunosuppressive activity with the goal of intercepting PDAC development.

ST analysis of the PanINs also identified transcriptional signatures that are known to be associated with PDAC phenotypes. PDACs are classified into classical and basal-like transcriptional subtypes^20^. Classical PDACs present a better prognosis and represent most tumor cells found in early stage cancers before patients receive treatment. This supports the hypothesis that initially all PDACs develop from the classical phenotype and that during the tumorigenesis there is a diverging point in which some cells will differentiate into the basal-like phenotype, a phenotype that is usually expanded by chemotherapy as resistance develops^20,36^. Further supporting this hypothesis, we detected the classical signature in six out of seven PanINs. The only sample that could not be classified as classical or basal-like expressed a CSC signature.

CSCs drive aggressive disease and their presence is associated with resistance to therapies, local recurrence, and development of metastasis^37–39^. The presence of CSCs and CAFs in PanINs suggest that the features associated with resistance to therapies in PDAC arise early during tumorigenesis. Further studies are necessary to determine how the interactions between these cell types can modulate additional features of resistance to therapies and progression of PDACs.

The differential expression analysis shows that LG and HG PanINs are transcriptionally similar. Among the few genes differentially expressed between these two PanIN grades, *TFF1*, frequently up-regulated in PanIN and PDAC, demonstrated gradual increase during PanIN progression but little is known about its role in tumorigenesis. As mentioned previously, secreted TFF1 could be involved in tumor cell interactions with CAFs^27,40,41^ The concomitant presence of CAFs and high levels of *TFF1* should be further investigated to pinpoint the relationship between *TFF1* expressing neoplastic cells and the adjacent CAFs to determine the involvement of this gene in the CAF-PDAC cross-talk.

The computational approach we developed to analyze PanIN is novel and broadly applicable for multi-omic studies of carcinogenesis. While imaging data has been shown to refine clusters in ST analysis^14,60^, this is the first study to our knowledge to use large scale databases of imaging data to enable a machine-learning based annotation of broad cell subtypes at a single-cell resolution^16^. This approach is uniquely enabled by the FFPE extension of the ST technology, as FFPE based datasets are necessary to preserve cellular morphologies to leverage databases of pathology annotations. Moreover, the reliance on this approach on cellular morphology also enables potential future extensions to other imaging-based cellular classifiers of cellular function^61^. This imaging-based data integration approach (ST and CODA) enables accurate and automated identification of the transcriptional profiles of neoplastic cells across stages of PanIN development through the differential expression analysis described above. However, fully relating these neoplastic cell state transitions to cancer progression requires relating these transcriptional states across the transition from normal epithelial through pre-malignancy to malignant PDAC cells. Here, we demonstrate that transfer learning approaches developed to integrate scRNA-seq data^5,13,19^ with matched imaging data can be extended to relate spatial data from precancer to reference tumor atlases. In this study, this two-stage computational approach integrating imaging, ST, and scRNA-seq data enabled us to identify a transition from inflammatory signaling in neoplastic cells from low grade PanIN to cellular proliferation in later stages of carcinogensis. In our complementary atlas study that identifies this inflammatory signaling, we further correlated this transition with CAF abundance and validated the ability of CAFS to promote this signaling in a novel human organoid co-culture model. The downregulation of this inflammatory signaling through LG to HG PanIN lesions observed in the ST data in this current study may be due to development of immunosuppressive CAFs that are already present in PanINs and are functionally similar to those present in PDACs, as we demonstrated using a human co-culture organoid model^18^. In addition, CAFs can create an immunosuppressive TME by producing and releasing cytokines that inhibit immune cell infiltration and factors that provide cancer cells additional growth and survival advantages at later stages^42,43^. This hypothesis is further supported by our observation of enhanced increased growth pathways in the transition from LG to HG PanIN lesions observed in our study.

Although we used a limited sample number, we were able to corroborate previous findings related to PanINs and discover new features and their potential role in the progression of these lesions to a large-scale cohort of scRNA-seq data for invasive PDAC. We successfully visualized the microenvironment in which PanINs developed and showed for the first time the presence of CAFs with potential suppressive function in PanINs. Our cohort included samples with varying stromal and acinar cell composition, but due to the limited size we did not observe correlations between PanIN transcriptional profiles with the adjacent cell types. To examine if the CAFs surrounding the PanINs remodel the premalignant microenvironment and influence premalignancy progression, a larger cohort with a more stringent selection criteria that includes patients’ clinical features and outcomes (e.g.: tumor stage, metastasis, response to therapies) would be better suited to unveil the critical features of CAF-PanIN (or PDAC) interactions in PDAC tumorigenesis through correlations between clinico-pathological features and TME compostion. Nonetheless, we demonstrate that multi-omics analysis enabled by FFPE ST, imaging data, and scRNA-seq data lead to a model in which neoplastic cells transition from CAF-induced inflammation to cellular proliferation during PDAC carcinogenesis. Moreover, this hybrid experimental and computational approach for provides broadly applicable tools to create a molecular and cellular model of the pathways that underlie carcinogenesis from multi-modal data spanning distinct high-dimensional transcriptomics and spatial molecular technologies.

## Data and code availability

Submission of spatial transcriptomics data to dbGAP and code to github are in process.

## METHODS

### Sample selection

FFPE pancreatic ductal adenocarcinoma (PDAC) surgical specimens collected from 2016 to 2018 were examined by experienced pathologists (KF and LDW). PanINs present in the specimens were marked and selected for ST analysis and were classified as low-and high-grade by experienced pathologists (L.D.W. and K.F.). The samples were obtained from the Johns Hopkins University School of Medicine Department of Pathology archives under Institutional Review Board approval (IRB00274690) under a waiver of consent.

### RNA quality control

All samples selected for the study had their RNA quality checked prior to the ST slides preparation. Total RNA was isolated from 20um sections of each sample using the RNase FFPE kit (Qiagen), following manufacturer’s instructions. RNA quality was measured using the DV200 assay on the Bionalyzer (Agilent) to determine the proportion of fragments with ~200bp in the sample. RNA quality was considered good if DV200 > 50%.

### Spatial transcriptomics slide preparation

The ST data was generated using the commercial platform Visium FFPE (10x Genomics). The slides are designed to accommodate a total of 4 sections with a maximum size of 6 x 6 mm. For the specimens that were larger than the designated regions of the Visium slides, we scored the selected sample area containing the PanIN using skin punches of 5mm in diameter. The skin punches were used directly on the FFPE blocks to delimit the area of interest, so when the block was sectioned in the microtome the PanIN containing region was detached from the rest of the section and could then be placed in the ST capture area of the slides (**Figure 1A**). A 5μm section from each sample with 5mm in diameter was used for the ST analysis. Upon preparation, the slides were incubated at 42°C and then stored in a desiccator until use.

### Spatial transcriptomics data generation

Using the Visium FFPE (10x Genomics) platform and following manufacturer’s validated protocol the samples were deparaffinized, stained with hematoxylin, and scanned using the Nanozoomer scanner (Hamamatsu) at 40x magnification. Human probe hybridization was performed overnight at 50°C. Following probe ligation, the RNA was digested, and the tissue was permeabilized for the release, capture, and extension of the probes. The designated area for each sample is covered by probes containing oligo-d(T) that capture the probes by a poly-A tail sequence present in the probe sequence. The sequencing library preparations were performed as instructed by the manufacturer using the extended probes as the template. All libraries were sequenced with a depth of at least 50,000 reads per spot (minimum of ~250 millions per sample) at the NovaSeq (Illumina). The Visium Human Transcriptome Probe Set v1.0 contains probes to 19,144 genes and after computational preprocessing (filtering for probes off-target activity) provides gene expression information for 17,943 genes.

### Cell type annotation using transfer learning from H&E imaging

Seven microanatomical components of human pancreas tissue were multi-labelled using a semantic segmentation workflow. The seven components recognized were (1) islets of Langerhans, (2) normal ductal epithelium, (3) vasculature, (4) fat, (5) acinar tissue, (6) collagen, and (7) pancreatic intraepithelial neoplasia (PanIN). Briefly, fifty examples of each tissue type were manually annotated using Aperio ImageScope. Half of the newly generated annotations were used in the training dataset for the convolutional neural network and the other half were used as an independent testing dataset to evaluate model performance. The testing dataset revealed an overall accuracy of 94.0% in classification of tissues in the TMAs. Following training, the tissue images were segmented to a resolution of 1μm.

Nuclear coordinates were generated via detection of two-dimensional hematoxylin intensity peaks. Briefly, the TMA images were downsampled to a resolution of 1 μm/pixel. As the tissues contained only a hematoxylin signal, color deconvolution (generally used to de-mix the hematoxylin channel from the hematoxylin & eosin image) was not necessary. Instead, the color image was converted to greyscale. The image was smoothed using a Gaussian filter and twodimensional intensity peaks with minimum radii of 2μm were identified as nuclear coordinates.

### Registration of ST data with cell type annotations

The low-resolution image used for the Visium pre-processing with Space Ranger was registered to the high-resolution tissue image used for microanatomical measurements to integrate the two workflows. The registration utilized the fiducial markers present on the ST glass slide to estimate the registration scale factor and translation. As registration was performed on two scans of identical tissue sections, it was assumed that rotation was not necessary. Here, the low-resolution image was registered to the high-resolution image (rather than the other way round) so that the scale factor was always greater than 1 and ensuring that the 1 μm resolution of the tissue micro annotations was preserved. First, the fiducial markers in each pair of images were segmented by identification of small, nonwhite objects surrounding the larger TMAs. Nonwhite objects were determined to be pixels with red-green-blue standard deviations greater than 6 in 8-bit space. These objects were morphologically closed and very small noise (<50 pixels) were removed. The fiducial markers were then determined to be objects in the image within 20% of the median object size (as many fiducial markers existed for each corresponding tissue image). This process resulted in fiducial image masks for the high-resolution and low-resolution tissue images. With these masks, four possible registrations were calculated to account for the situation where the Visium analysis was performed on the tissue image rotated at a 0-, 90-, 180-, or 270-degree angle. For each registration, the corner fiducial markers of the low-resolution image were rescaled and translated to minimize the Euclidean distance to the fiducial markers of the high-resolution image. Of the four registration results, the registration resulting in the greatest Jaccard coefficient between the high-resolution and low-resolution fiducial masks was chosen. For the eight TMAs, the average Jaccard coefficient of the fiducial masks was 0.94.

The registration information used to overlay the low-resolution tissue image to the high-resolution tissue image was used to convert the coordinates corresponding to the location of the Visium assessment in the low-resolution image into the high-resolution images coordinate system. Once the Visium coordinates were registered to the high-resolution image, the generated tissue microanatomy composition and cellularity were calculated for regions within 25μm of each coordinate. For each Visium coordinate, pixels in the micro-anatomically labelled mask image within 25μm of that coordinate were extracted. Tissue composition was determined by analyzing the % of each classified tissue type within that dot. The cellularity of each dot was determined by counting the number of nuclear coordinates within 25μm of each Visium coordinate. Cellular identity was estimated by determining the microanatomical label at each coordinate where a nucleus was detected (a nucleus detected in the same pixel where the semantic segmentation model detected normal ductal epithelium was labelled an epithelial cell).

### Spatial transcriptomics data analysis of PanIN samples

Sequencing data was processed using the Space Ranger software (10x Genomics) for demultiplexing and FASTQ conversion of barcodes and reads data, alignment of barcodes to the stained tissue image, and generation of read counts matrices. The processed sequencing data were inputs for the analyses using the Seurat software^1–4^. Data preprocessing with Seurat involved initial visualization of the counts onto the tissue image to discriminate technical variance from histological variance (e.g.: collagen enriched regions present lower cellularity that reflects in low counts). The filtered data was normalized using the SCTransform approach that uses a negative binomial method to preserve biological relevant changes while filtering out technical artifacts. Following normalization, data from all slides were merged and batch correction was performed with Harmony from harmony_0.1.0. Unsupervised clustering was subsequently performed on the harmony reduction using the Louvain algorithm as implemented by Seurat.

Louvain clusters were annotated using the overlap of CODA annotations and quantifications per spot with well-characterized marker genes. Neoplastic and ductal epithelium groups were generated through selecting spots from the respective Louvain cluster that were estimated to be greater than or equal to 70% of the respective cell type on CODA. The data dimensionality was reduced using PCA for clustering and in tissue visualization of the transcriptional clusters. Unsupervised clustering was performed based on the most variable features (genes). Differential gene expression analysis of normal ducts and PanINs, and low and high grade lesions were performed using the MAST test^5^ as implemented by Seurat. For comparisons performed across different slides, the slide was assigned as a latent variable and the matrix was prepared using *PrepSCTFindMarkers* to account for the multiple SCT models. Pathway analysis was performed using GSEA v4.2.1^6,7^. High-and low-grade PanIN spots were subset from the neoplastic Louvain cluster by pathologist (LDW) annotation using a custom-made Shiny app derived from the *SpatialDimPlot* function in Seurat.Violin plots, spatial plots, were generated in Seurat. Volcano plots were generated in ggplot28. Heatmaps were generated using ComplexHeatmap9.

### Transfer learning to relate ST data from PanIN to a scRNA-seq atlas of Pancreatic Ductal Adenocarcinoma

We obtained scRNA-seq data for pancreatic epithelial cells from an atlas of 29 tumor samples and 14 non-cancerous samples collated from Peng et al. and Steele et al. as described in Kinny-Koster et al.^10^. We inferred cellular phenotypes in the epithelial cells using CoGAPS (R, version 3.5.8)^11,12^ to infer 8 patterns on the log transformed expression values. Pattern annotation was based on overrepresentation analysis of patternMarker genes identified by CoGAPS (R, version 3.9.5)^13^ and Molecular Signatures Database Hallmark gene sets (version 7.5.1)^14,15^ using the R package fgsea (version 1.18.0)^16^. *TFF1* expression was measured as log-normalized counts. Uniform manifold approximation and projection (UMAP) plots were made using monocle3 (version 1.0.0)17–23. UMAP plots for epithelial cells from the PDAC atlas were made with cells colored by epithelial cell type, log normalized *TFF1* expression, and Pattern 2, 5, 7 weights.

PanIN ST data was subset to spots annotated as epithelial by CODA (N = 623 spots; normal = 254, low-grade = 110, high-grade = 159). CoGAPS patterns learned from the PDAC atlas were projected onto scaled SCT expression values from epithelial ST spots using ProjectR (version 1.8.0)^24,25^. Projected pattern weights were plotted as violin plots using Seurat (version 4.1.0). Mean pattern weights were compared across epithelial lesion grades using Wilcoxon rank-sum tests within ggpubr (version 0.4.0). UMAP plots of ST spots and over layed plots of ST spots colored by epithelial type, log normalized *TFF1* expression, and projected Pattern 2, 5, 7 weights over tissue slices were prepared using Seurat (version 4.1.0)^1^.

## Data and code availability

Submission of spatial transcriptomics data to dbGAP and code to github are in process.

## Acknowledgements

The authors would like to thank the Oncology Tissue Services (OTS) Core Facility and the Genetic Resources Core Facility (GRCF) for tissue sections preparation and library sequencing services, respectively. This work was supported by The Sol Goldman Pancreatic Cancer Research Center grant (to L.T.K.), NIH P01-CA247886-01A1 (to E.M.J., E.J.F and L.T.K), SU2C/AACR DT-14-14 (to E.M.J.), Lustgarten Foundation Pancreatic Cancer Research grant (to E.M.J., E.J.F and L.T.K), the Emerson Cancer Research Fund (to E.M.J., E.J.F.), an Allegheny Health Network (AHN) grant (to E.J.F.), NIH U01CA212007 (to E.J.F.), NIH U01CA253403 (to E.J.F.), the JHU Discovery Award (to E.J.F.), and SPORE GI P50CA062924-24A1 (to E.M.J, E.J.F. and L.T.K), NCI F31CA268724-01 (to D.N.S), NIH U54CA268083 (to D.W., P.W., A.L.K.), NIH U54CA210173 (D.W.), NIH U01AG060903 (to D.W.), Susan Wojcicki and Dennis Troper (to A.L.K.), The Rolfe Foundation for Pancreatic Cancer Research (to A.L.K.)

## Competing Interest Declaration

E.M.J. reports other support from Abmeta, personal fees from Genocea, personal fees from Achilles, personal fees from DragonFly, personal fees from Candel Therapeutics, other support from the Parker Institute, grants and other support from Lustgarten, personal fees from Carta, grants and other support from Genentech, grants and other support from AstraZeneca, personal fees from NextCure and grants and other support from Break Through Cancer outside of the submitted work. E.J.F. is on the Scientific Advisory Board of Viosera Therapeutics/Resistance Bio and is a consultant to Mestag Therapeutics. No disclosures were reported by the other authors.

## Author’s Contributions

*A.T.F. Bell, J.T. Mitchell, A.L. Kiemen:* data curation, formal analysis, investigation, visualization, writing original draft and review. *K. Fujikura:* data curation, writing and review. *H. Fedor, B. Gambichler, A. Deshpande, P.H. Wu, D.N. Sidiropoulos, R. Erbe, R. Chan, S. Williams, J. Chell, J. Zimmerman., D. Wirtz:* resources, writing review. *E.M. Jaffee:* resources, writing review and editing. *L.D. Wood:* data curation, resources, writing review and editing. *E.J. Fertig:* conceptualization, data curation, supervision, formal analysis, visualization, writing original draft, review and editing. *L.T. Kagohara:* conceptualization, data curation, supervision, formal analysis, funding acquisition, visualization, writing original draft, review and editing.

## Supplementary Information

Supplementary Information is available for this paper

## Notes

### Summary of Updates

Figures were revised to add a panel of the analysis performed to generate the results for clarity. The abstract, introduction and discussion were updated to highlight the novel approaches used.

