## Supplemental figures for "PanIN and CAF Transitions in Pancreatic Carcinogenesis Revealed with Spatial Data Integration"

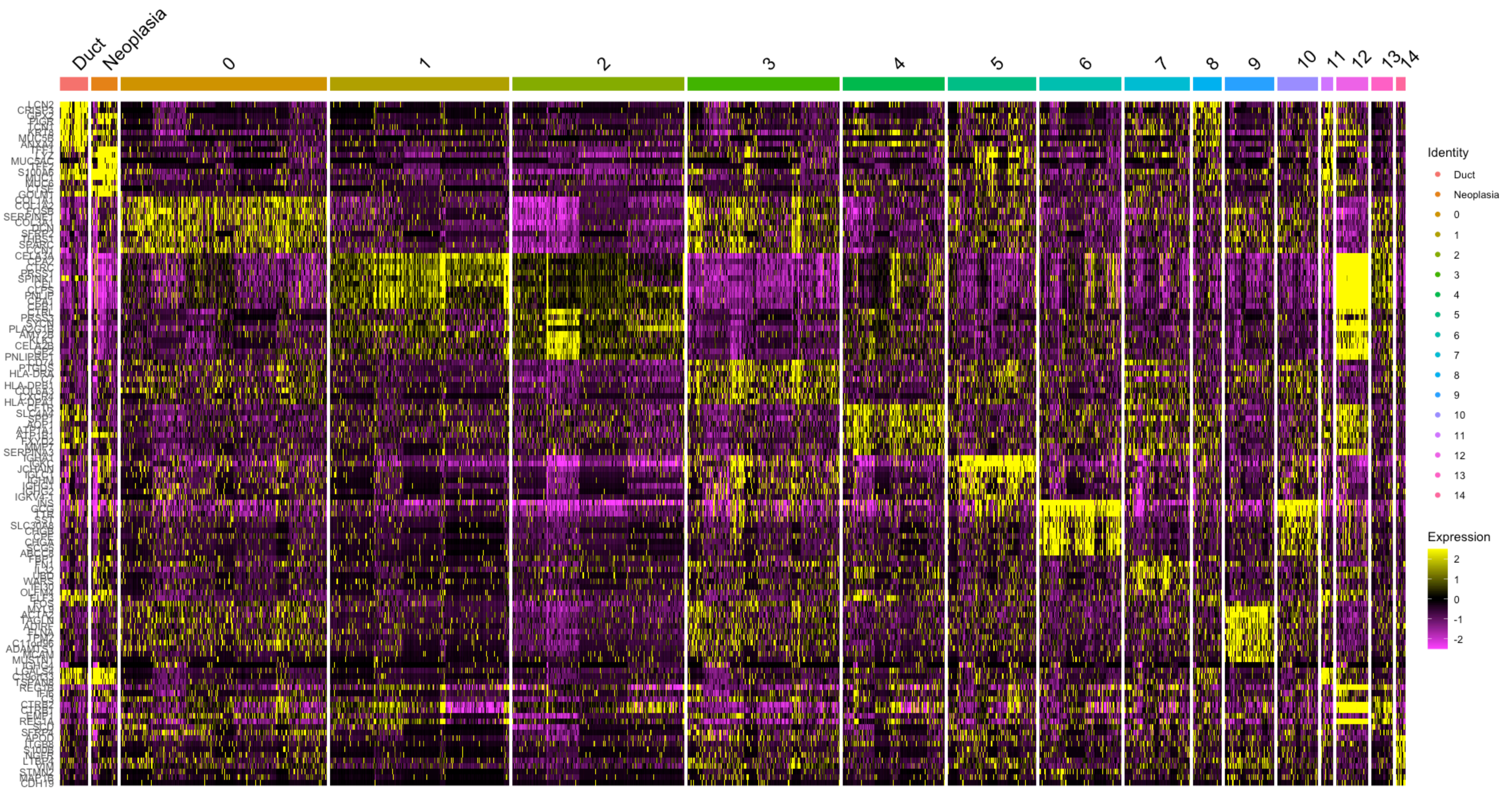


Supplemental Figure 1 - Heatmap representing the expression of genes used to identify the clusters in the spatial transcriptomics data.

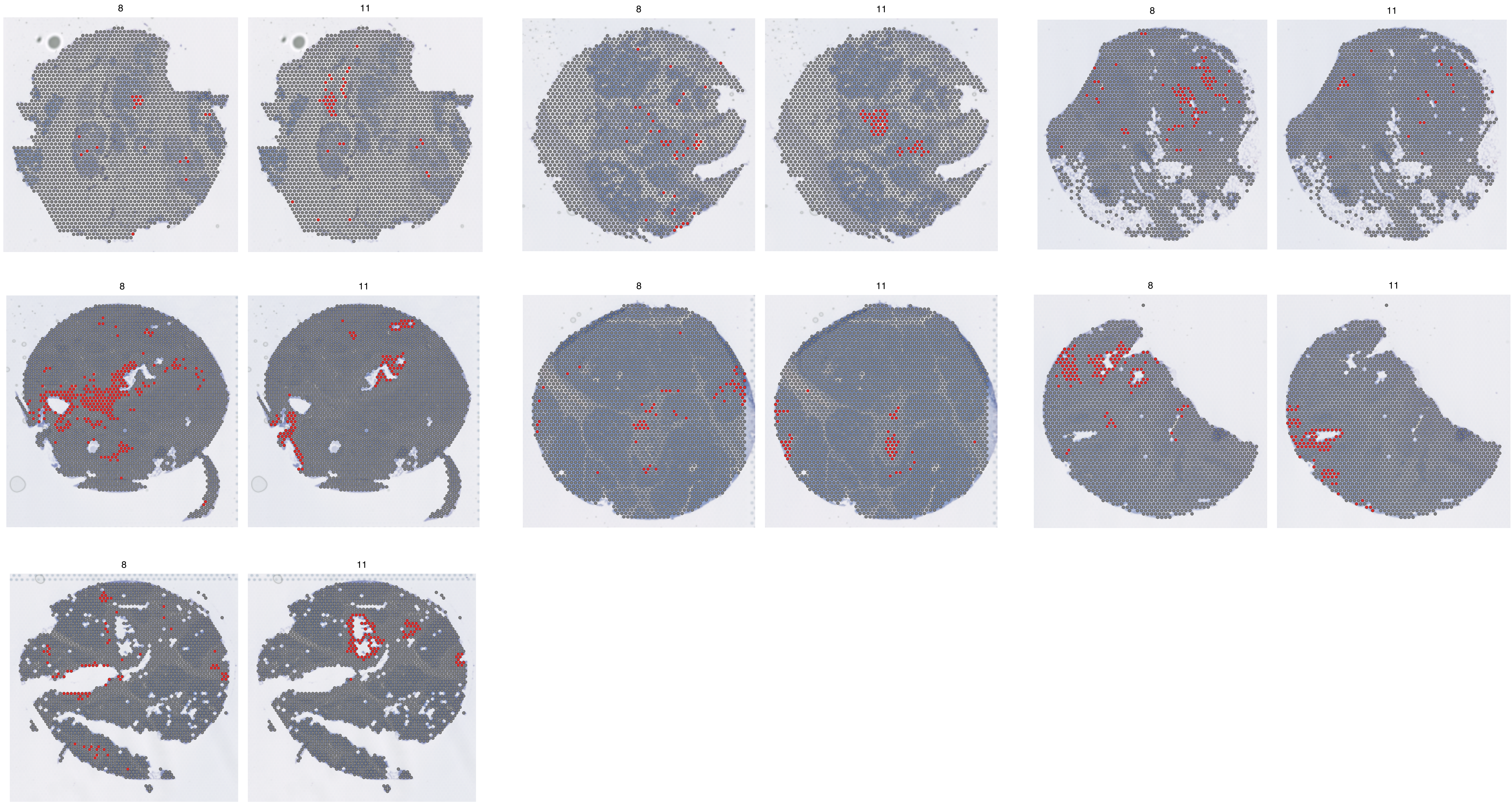


Supplemental Figure 2 - Representation of normal ducts (cluster #8) and pancreatic intraductal neoplasia (PanIN) (cluster #11) clusters identified by unsupervised clustering using Seurat.

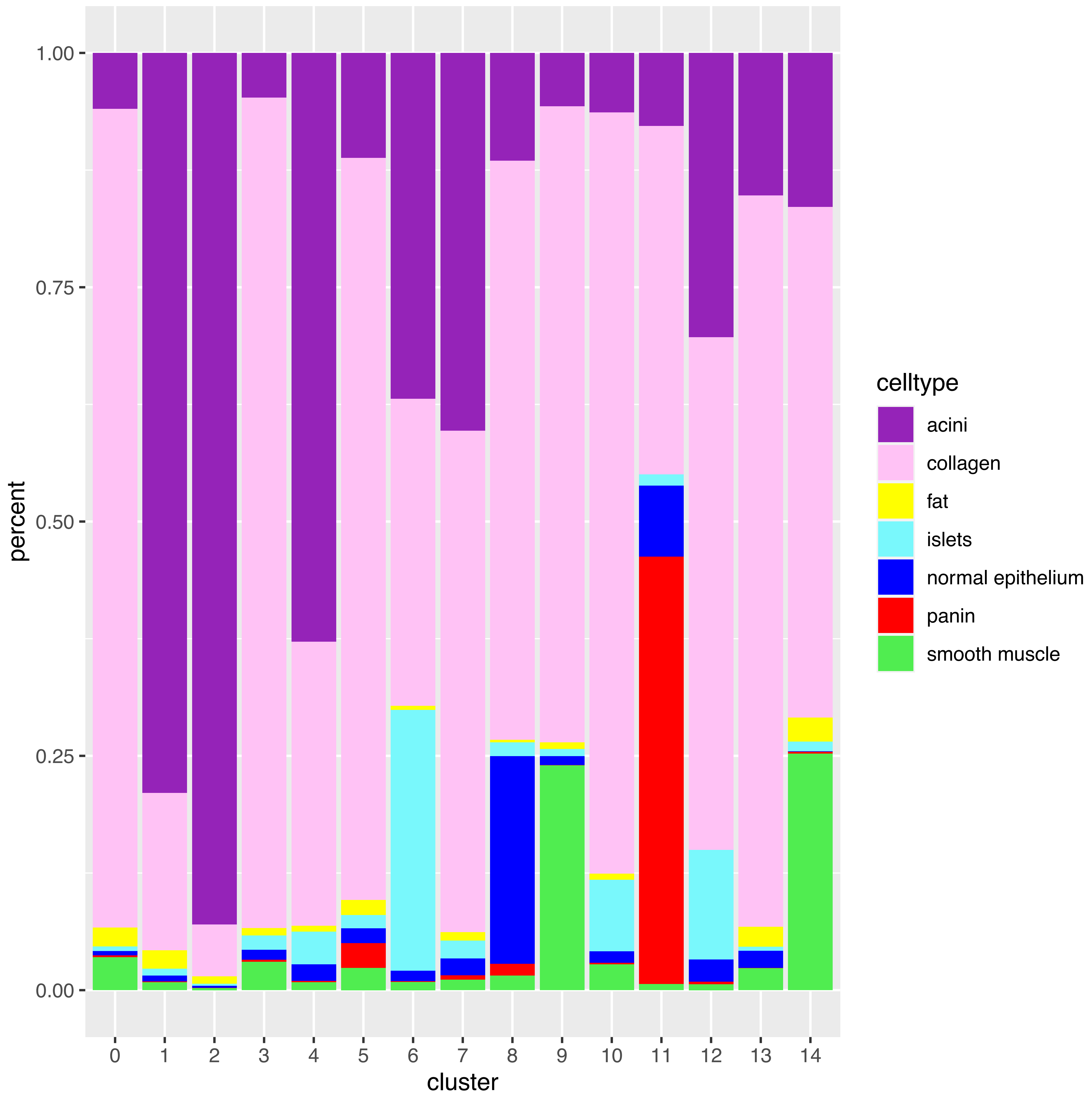


Supplemental Figure 3 - Cell type proportions annotated in each gene expression cluster using transfer learning (CODA). Clusters encompassing spots with similar gene expression profiles, included more than one cell type due to the lack of single-cell resolution inherent of ST approach. CODA identifies cell types learnt from their morphology and was further used to precisely select spots encompassing a single cell type.

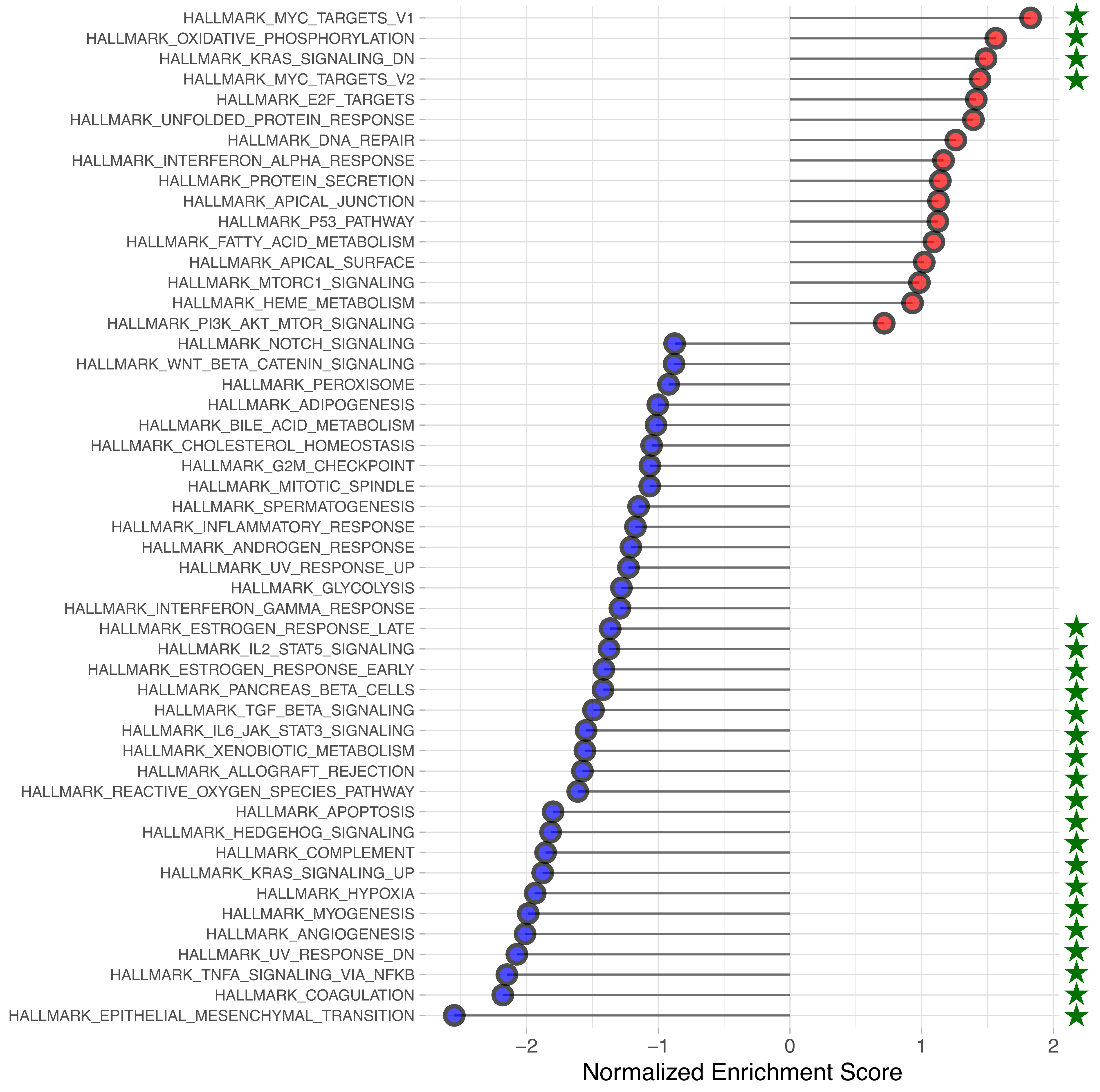


Supplemental Figure 4 - Gene set enrichment analysis (GSEA) of the differentially expressed genes between PanIN versus normal ducts. Red circles mark pathways enriched in PanINs and blue circles mark pathways enriched in normal ducts. The significantly enriched pathways are marked with green stars.

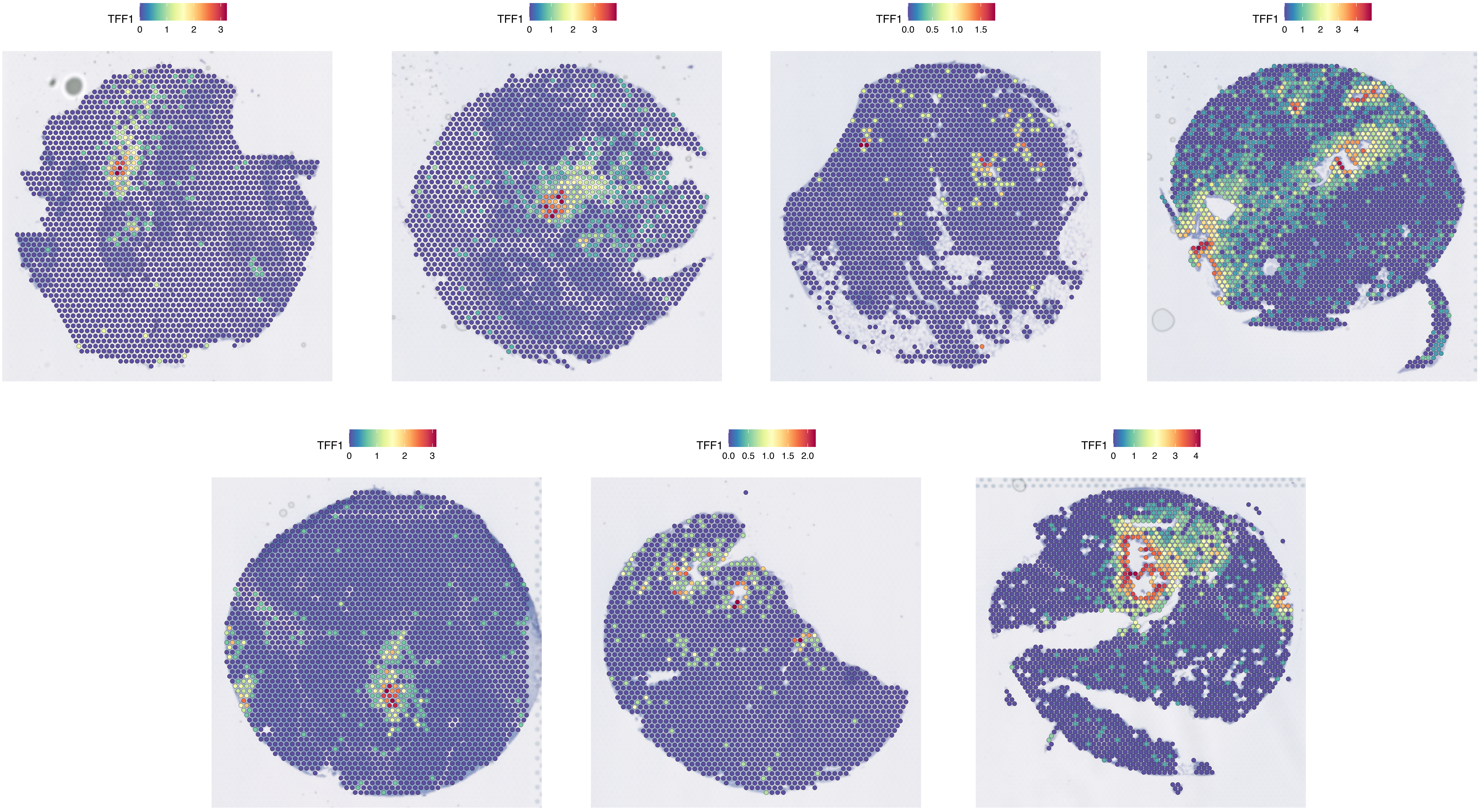


Supplemental Figure 5 - TFF1 expression. TFF1 is highly expressed in PanINs (red dots) in six out of seven samples. The only PanIN that does not express TFF1 is the lesion that expresses the cancer stem cell signature (middle bottom panel).

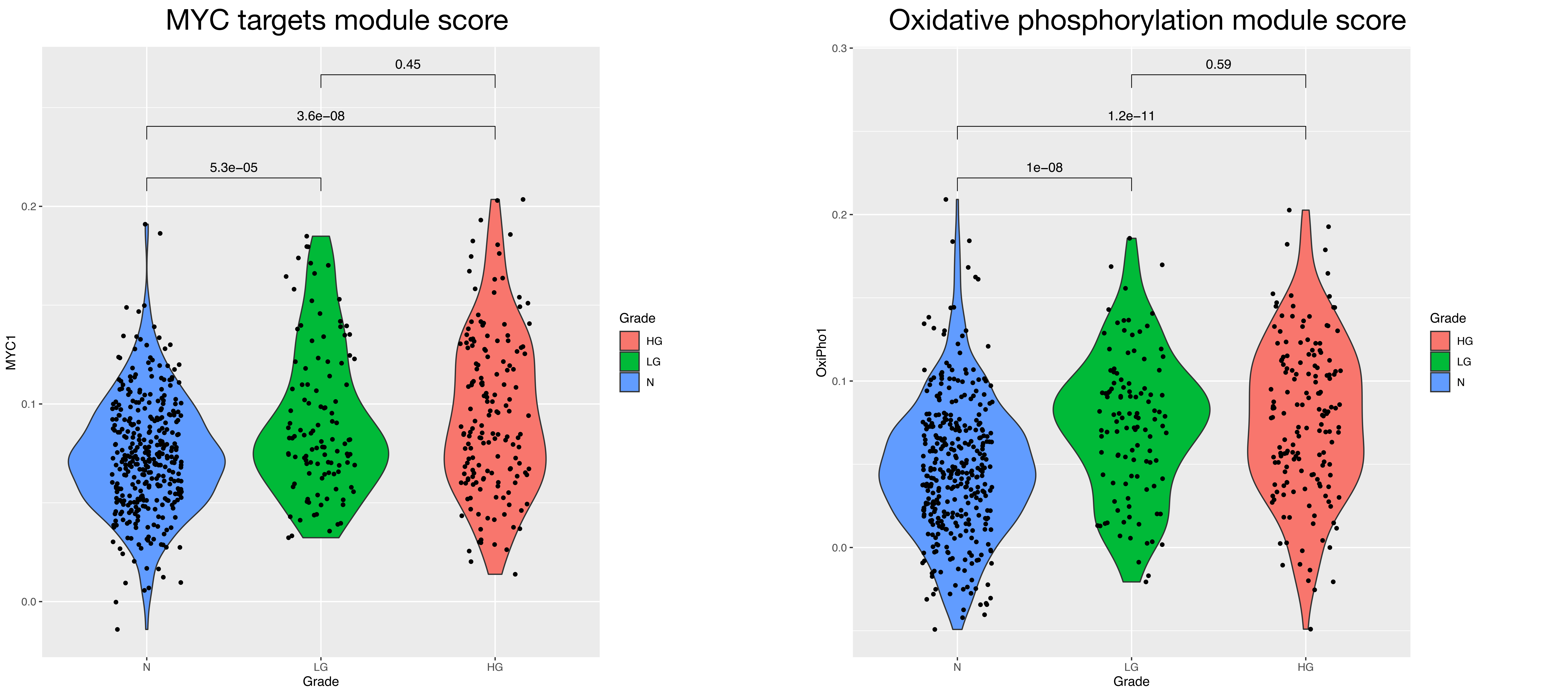


Supplemental Figure 6 - Violin plots representing MYC targets and oxidative phosphorylation (OXPHOS) pathway module scores. Module scores for MC targets and OXPHOS were calculated using the Seurat function AddModuleScore( ), using the list of genes from the MSigDB Hallmark pathways (HALLMARK_MYC_TARGETS_V1, HALLMARK_MYC_TARGETS_V2 and HALLMARK_OXIDATIVE_PHOSPHORYLATION). For MYC, the HALLMARK_MYC_TARGETS_V1 and HALLMARK_MYC_TARGETS_V2 were merged. MYC targets and OXPHOS pathways were enriched in the differential expression analysis comparing PanINs versus normal ducts (N). Those pathways are up-regulated in PanINs but there is no significant difference in expression between low (LG) and high grade (HG) pre-malignant lesions.

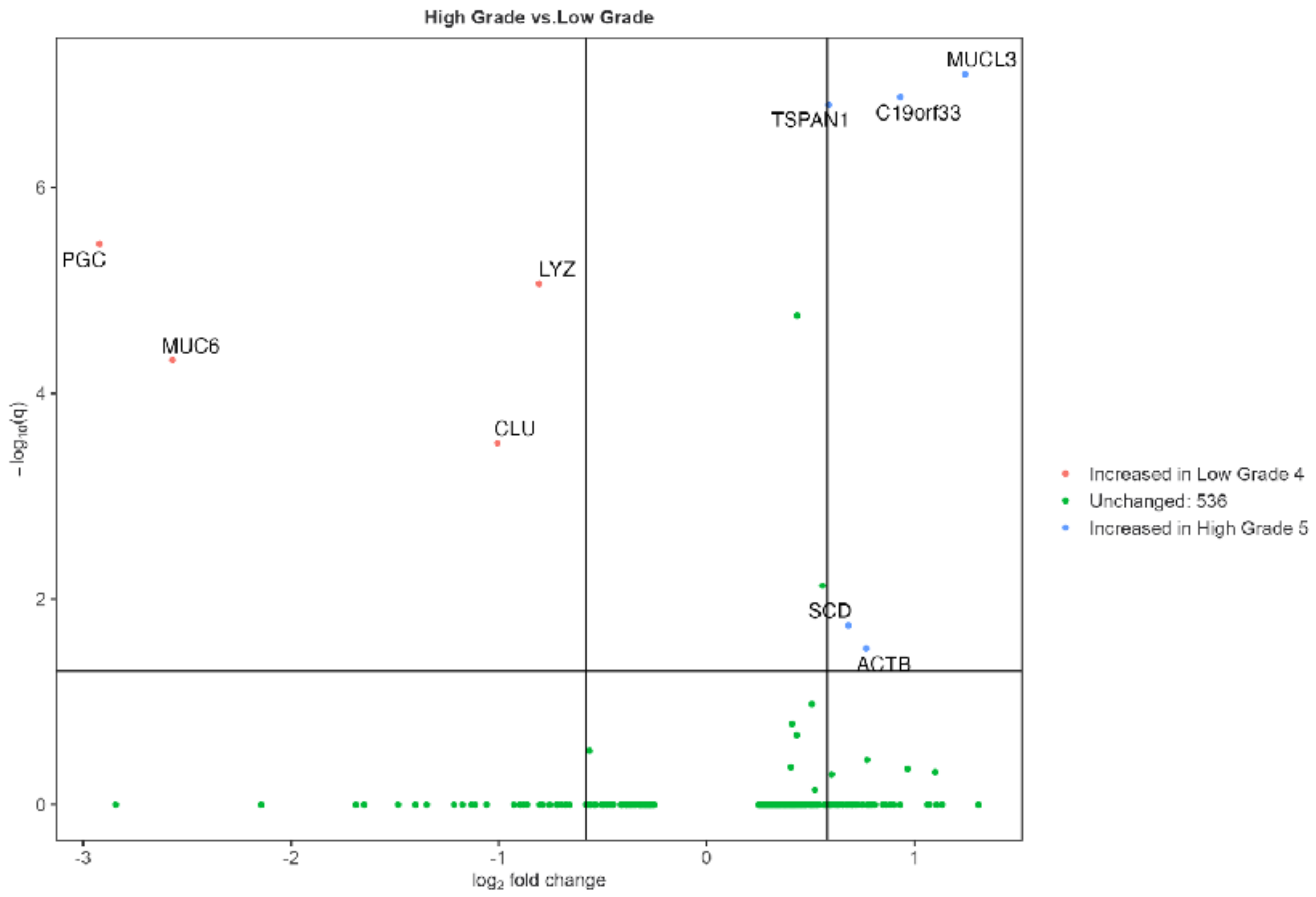


Supplemental Figure 7 - Volcano plot of the differential expression analysis (DEA) of high grade (HG) relative to low grade (LG) PanINs. The DEA identified 5 genes significantly up-regulated in HG lesions (blue) and 4 genes up-regulated in LG lesions (red).

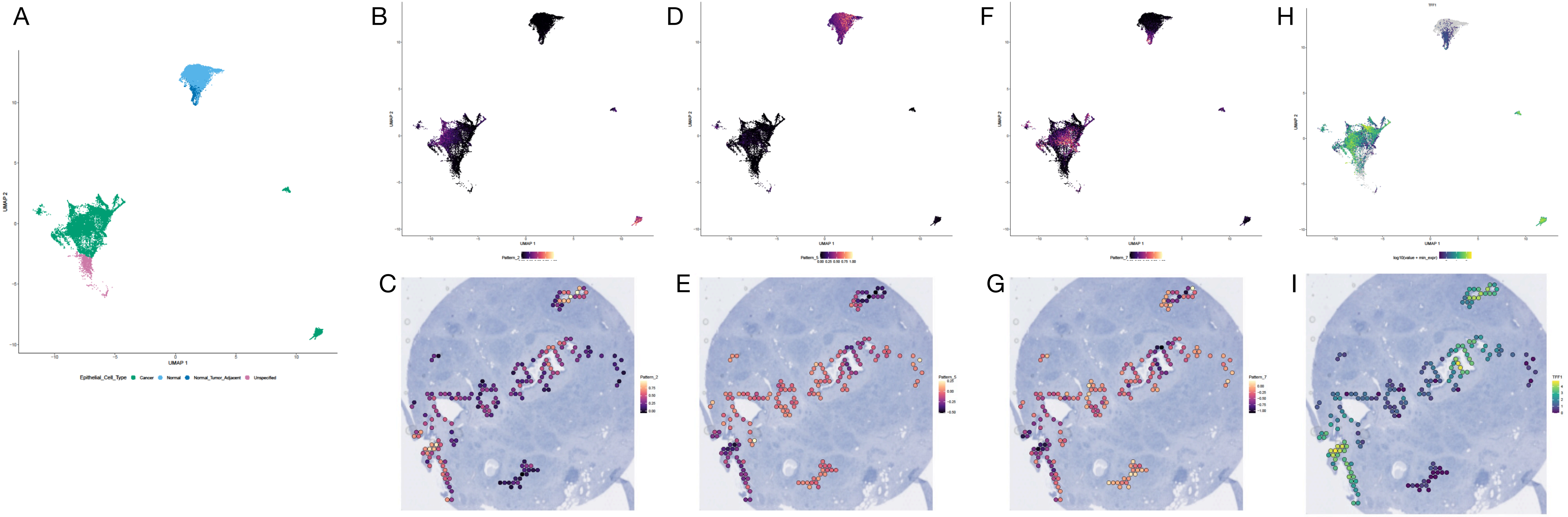


Supplemental Figure 8 - PDAC atlas and PanIN ST transfer learning. A. UMAP of tumor and normal epithelial cells from the PDAC atlas. B. Pattern 2 weights enrichment on PDAC atlas epithelial cells. C. Representation of Pattern 2 weights enrichment on PanIN ST data of a mixed duct. D. Pattern 5 weights enrichment on PDAC atlas epithelial cells. E. Representation of Pattern 5 weights enrichment on PanIN ST data of a mixed duct. F. Pattern 7 weights enrichment on PDAC atlas epithelial cells. G. Representation of Pattern 7 weights enrichment on PanIN ST data of a mixed duct. G. *TFF1* expression on PDAC atlas epithelial cells. H. *TFF1* expression on PanIN ST data of a mixed duct.
